# Humanized CAR T cells targeting p95HER2

**DOI:** 10.1101/2022.05.20.492812

**Authors:** Macarena Román, Irene Rius-Ruiz, Ariadna Grinyó-Escuer, Santiago Duro-Sánchez, Marta Escorihuela, Ekkehard Moessner, Christian Klein, Joaquín Arribas

**Author notes:** **Corresponding author:** Joaquín Arribas, PhD, Hospital del Mar Medical Research Institute (IMIM), C/ Doctor Aiguader 88, Barcelona 08003, Spain.

## Abstract

Redirection of T lymphocytes against tumor-associated or tumor-specific antigens, via bispecifc T cells engagers (BiTEs) or chimeric antigen receptors (CARs), is a successful therapeutic strategy against certain hematologic malignancies. In contrast, so far it has failed against solid tumors. Given the scarcity of tumor-specific antigens, the vast majority of BiTEs and CAR Ts developed to date have been directed against tumor-associated antigens. These are expressed in some normal tissues and, as a consequence, frequent and serious side effects caused by on-target off-tumor activity have limited the use of BiTEs and CARs in the clinic. P95HER2 is a fragment of the tyrosine kinase receptor HER2 expressed in more than 30% of HER2-amplified tumors. It has been previoulsy shown that p95HER2 is a tumor-specific antigen. Here we present the generation of CARs targeting p95HER2. p95HER2 CAR T cells show remarkable activity against p95HER2-expressing cells *in vitro* and *in vivo*. Further, they are also effective against lung and brain metastasis. These tumor-specific CAR T cells could be used in the near future to deliver additional anti-tumor therapies in a safe manner.

## Introduction

Redirection of T lymphocytes against tumor-associated or tumor-specific antigens is a successful therapeutic strategy against certain hematologic malignancies. Despite the promise of these therapies, so far, they have failed against solid tumors.

T cells can be redirected against cancer cells using bispecific antibodies, known as bispecific T cell engagers (BiTEs) or T cell bispecific antibodies (TCBs). BiTEs simultaneously bind the tumor antigen and an invariant subunit of the T cell receptor (TCR), typically CD3 (Ellerman, 2019). Alternatively, T cells can be manipulated to express chimeric antigen receptors (CARs), which are engineered proteins containing the antigen-binding domain of an antibody and signaling domains of the CD3 subunit of the TCR as well as those of co-stimulatory receptors such as CD28 or 4-1BB (Waldman et al., 2020).

Given the scarcity of tumor-specific antigens, the vast majority of BiTEs and CAR Ts developed to date have been directed against tumor-associated antigens. These are expressed in some normal tissues and, as a consequence, frequent and serious side effects caused by on-target off-tumor activity have limited the use of BiTEs and CARs (June and Sadelain, 2018).

The gene encoding the tyrosine kinase receptor HER2 is amplified in approximately 3% of all tumors. In some tumors, such as those affecting the breast and the upper gastrointestinal tract, the frequency of HER2 amplification reaches ∼15%. More than one third of these tumors also express a truncated form of HER2 known as p95HER2 (Arribas et al., 2011). HER2 is also expressed in normal epithelium, albeit at much lower levels than in HER2-amplified tumors. In contrast, p95HER2 is a *bona fide* tumor-specific antigen (Ruiz et al., 2018). Confirming this observation, we have shown that a p95HER2 TCB effectively redirects lymphocytes against p95HER2-positive breast tumors while it has no effect on a variety of normal cells, some of them expressing HER2 (Ruiz et al., 2018).

Because of the therapeutic failures on solid tumors, efforts have focused on refining the strategies to redirect T cells. While improvement in the design of TCBs has inherent limitations, CAR Ts can be used as platforms to deliver additional anti-tumor factors such as cytokines or antibodies (June and Sadelain, 2018).

Here we present the development of CAR T cells targeting p95HER2. Characterization of these p95HER2 CAR Ts shows that they are remarkably effective against tumor cells expressing p95HER2, both in primary tumors and in metastases. Shortly, p95HER2 CAR Ts could be used to safely deliver additional anti-tumor therapies to p95HER2-expressing tumors.

## Results

### Generation and *in vitro* characterization of p95HER2 CAR Ts

The workflow to generate CAR Ts targeting p95HER2 is represented in Fig. 1. Briefly, out of 7 anti-p95HER2 monoclonal antibodies, we selected 3 because they stained cells expressing p95HER2 more intensely (Fig. 1B). The epitopes recognized by these anti-p95HER2 antibodies were very similar, but not identical. While the optimal epitope recognized by 14D and 15C antibodies is PIWKFPDE, the epitope optimally recognized by 32H2 is one amino acid shorter (Fig. 1C and (Ruiz et al., 2018)). The corresponding mouse sequence (PIWKYPDE), was not recognized by any of the anti-p95HER2 antibodies (Fig. 1D).

**Fig. 1.**
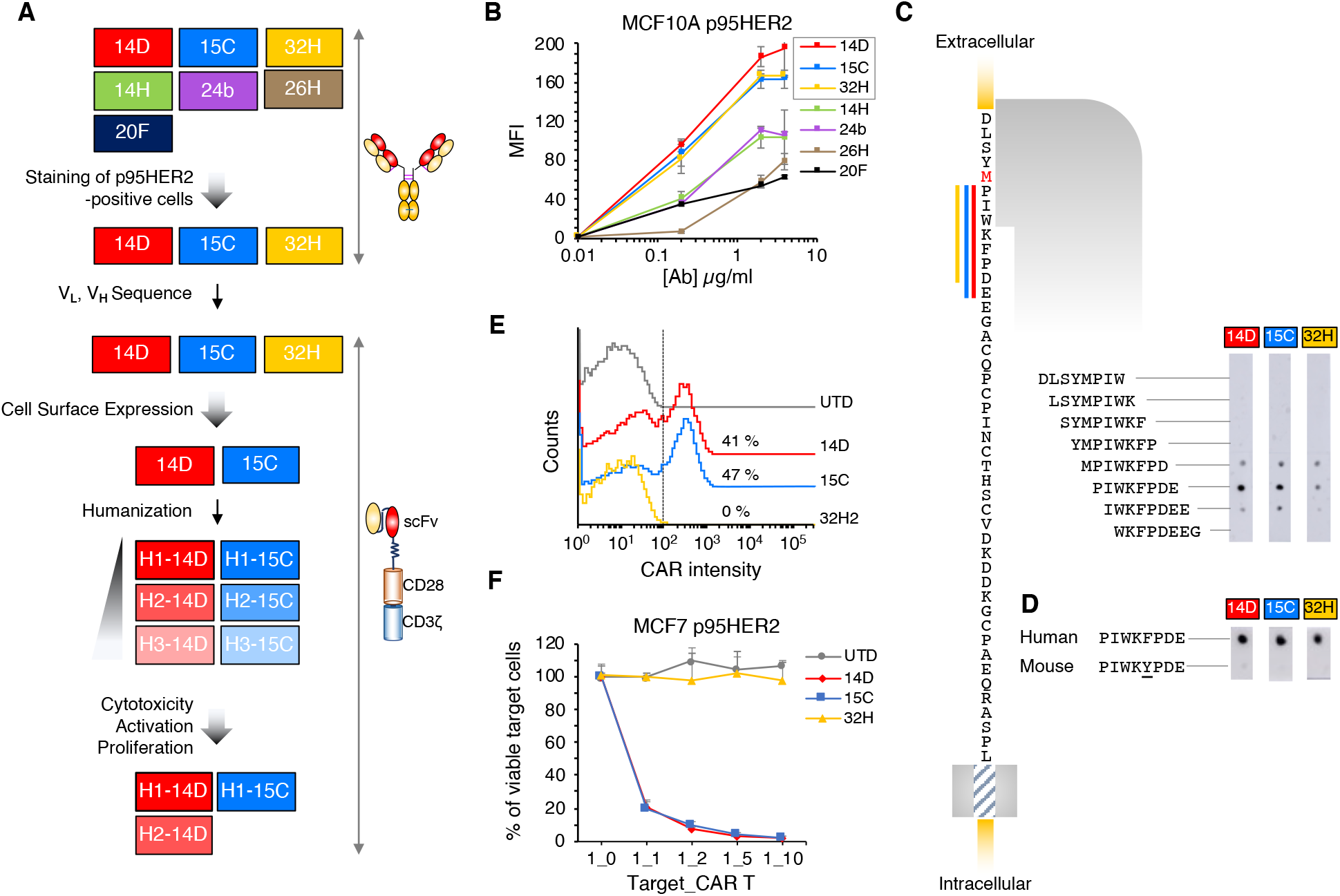
Generation and characterization of p95HER2-CARs. **A**, Schematic showing the workflow to generate p95HER2-CARs. **B**, MCF10A cells expressing p95HER2 (Parra-Palau et al., 2014) were stained with different concentrations of the indicated antibodies and analyzed by flow cytometry. Mean fluorescence intensities (MFI) are expressed as averages of four determinations. **C**, Schematic showing the amino acid sequence of the juxtamembrane extracellular region of HER2. p95HER2 starts at methionine 611 (highlighted in red). Overlapping 8-mer peptides were synthesized, immobilized and incubated with the indicated antibodies, an anti-mouse secondary antibody coupled to peroxidase and developed **D**, PIWKFPD and PIWKYPD peptides, corresponding to the human and mouse sequences, respectively, were synthesized, immobilized and incubated with anti-p95HER2s. Then, blots were processed as in C. **E**, The indicated CARs were transduced into PBMCs from healthy volunteer and their expression at the cell surface determined by flow cytometry. UTD, control non-targeting CAR. **F**, MCF7 cells expressing p95HER2 were co-cultured with different ratios of CAR T cells for 48 h. Then, viable target cells were quantified by flow cytometry using EpCAM as a marker. Results are expressed as averages of three independent experiments.

The sequences of the variable regions of the heavy and light chains of these antibodies were used to construct the scFv (single chain variable fragment) of second generation CARs containing CD28 and CD3z signaling domains. Only 2 of these CARs were expressed at the cell surface of T cells (Fig. 1E). Accordingly, only these two CARs exhibited cytotoxic activity against cells expressing p95HER2 (Fig. 1F).

The sequences of the murine scFv regions of these CARs were humanized to various degrees (Fig. 2). With the resulting sequences, we generated six humanized p95HER2 CAR Ts. Note that all the versions of the 14D CAR have the same variable light chain. The light and heavy chains corresponding to 15C were combined as indicated.

**Fig. 2.**
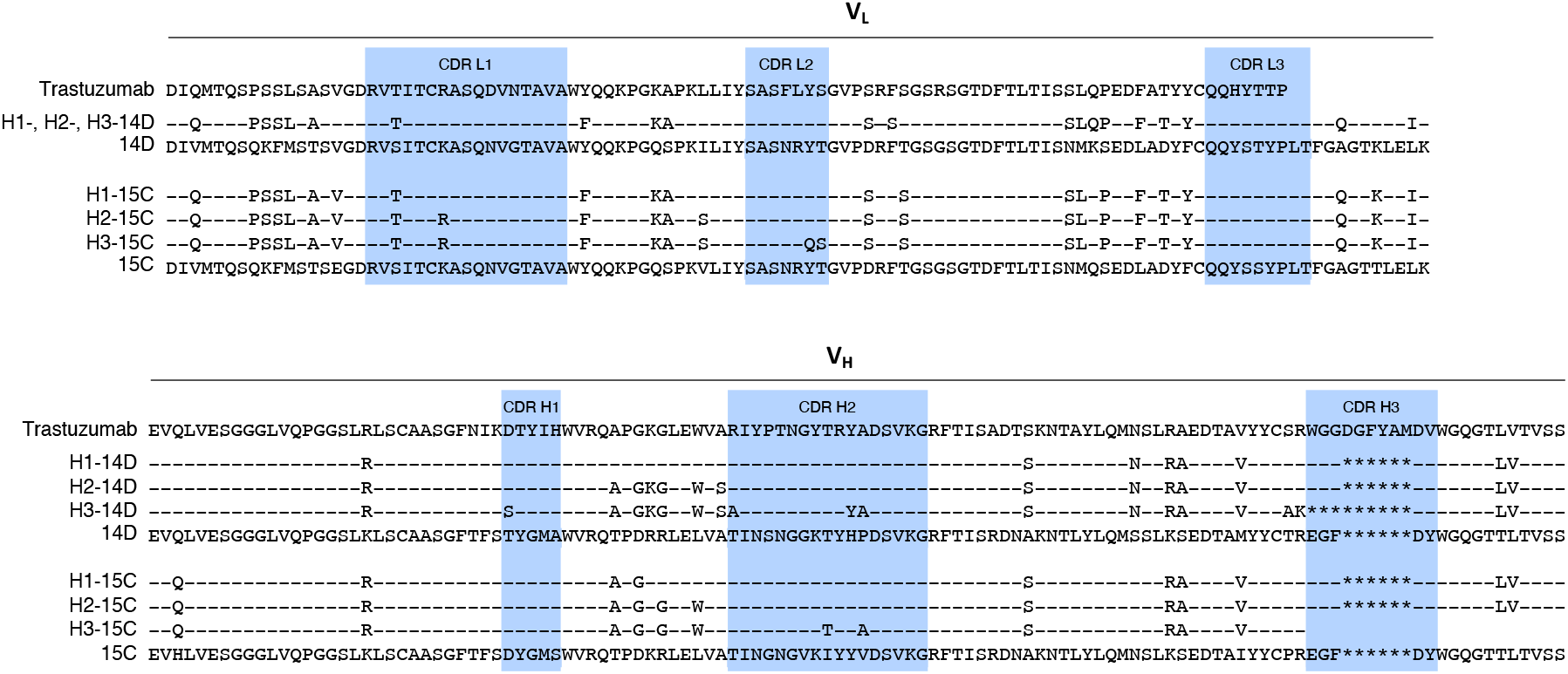
Humanization of the variable light (V_L_) and heavy (V_H_) chains of the 14D and 15C anti-p95HER2 antibodies. Using mRNAs from the corresponding hybridomas, the V_L_ and V_H_ chains were sequenced. The sequences of these antibodies humanized to various extents are shown. As references, we show the sequences of Trastuzumab and those of the 14D and 15C.

### *In vitro* efficacy of the humanized p95HER2 CAR Ts

The six humanized p95HER2 CAR Ts were characterized by co-culturing them with target cells expressing empty vector or p95HER2. In these co-cultures, CD8^+^ cells expressing three of the CARs tested (H1-14D, H2-14D and H1-15C) presented an statistically significant increase in the expression of the activation marker CD25 (Fig. 3A). Two of these CARs (H1-14D and H1-15C) promoted the proliferation of T cells (Fig. 3B). Despite increasing the expression of CD25, the H2-14D CAR did not promote T cell proliferation (Fig. 3B), confirming that the extent of activation induced by this CAR is comparatively lower. Supporting this conclusion, the cytotoxicity induced by the H1-14D and H1-15C CAR Ts was higher than that induced by H2-14D (Fig. 3C). Despite these differences, we selected the three p95HER2 CARs with detectable levels of activation and cytotoxic ability for further *in vivo* analyses.

**Fig. 3.**
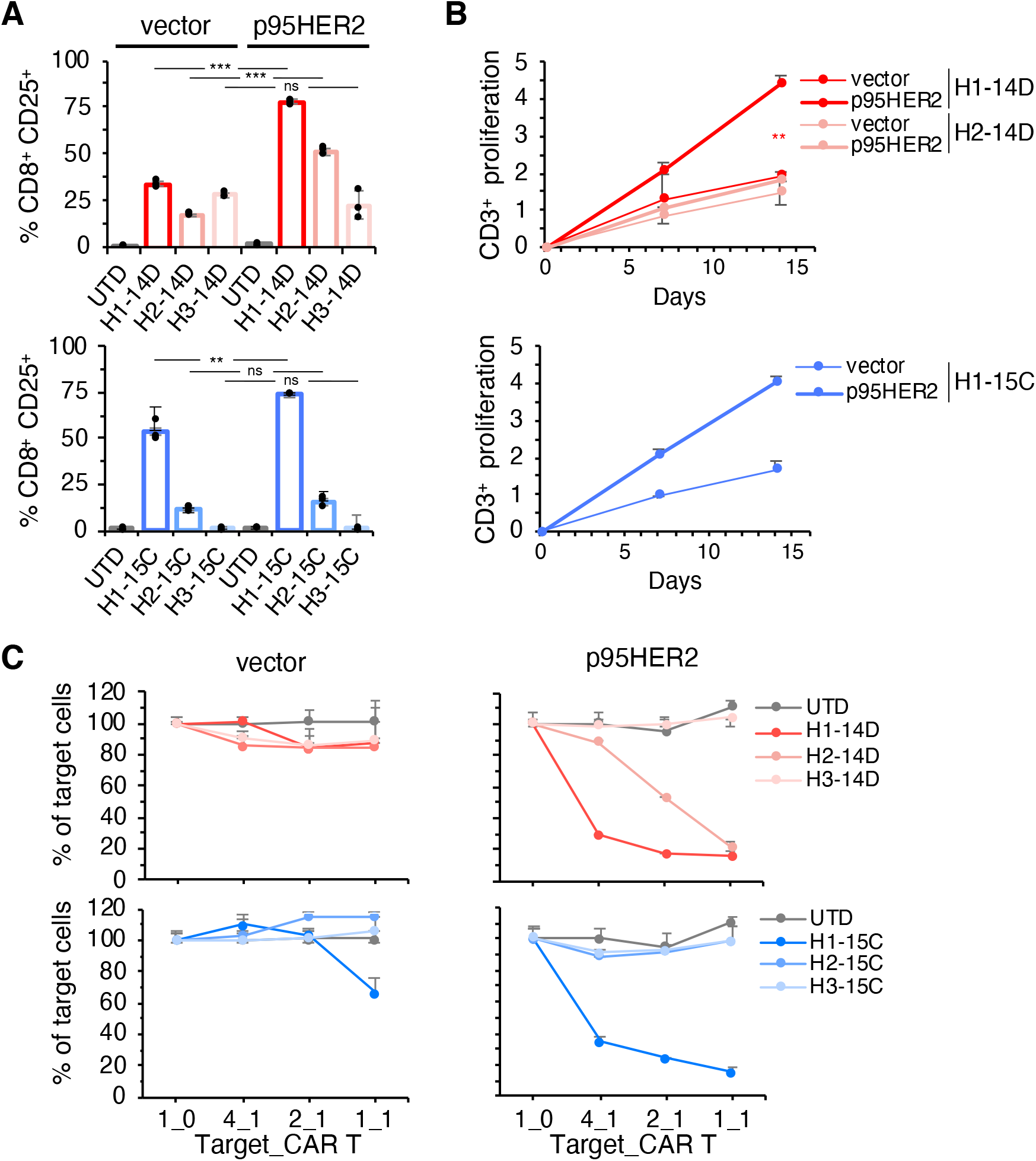
*In vitro* characterization of three humanized p95HER2 CAR Ts. **A**, MCF10A cells expressing empty vector or the same vector encoding p95HER2 were co-cultured CAR Ts at a ratio 2_1 for 48 h. Then, the numbers of CD8^+^CD25^+^ cells were determined by flow cytometry. Results are expressed as averages of three independent experiments. **B**, Numbers of CD3^+^ cells were determined by flow cytometry at different time points in the co-cultures described in A. Results are expressed as averages of two independent experiments. **C**, The same cells as in B, were co-cultured with different ratios of CAR T cells for 48 h. Then, viable target cells were quantified by flow cytometry using EpCAM as a marker. Results are expressed as averages of three independent experiments.

### *In vivo* efficacy of p95HER2 CAR Ts

The *in vivo* efficacy of the p95HER2 CAR H1-14D on MCF7 cells expressing p95HER2 was far superior than that of H1-15D (Fig. 4A). This superiority was not due to a higher infiltration of CARs into the tumor (Fig. 4B). Similarly, compared to that of H2-14D, the antitumor activity of H1-14D was higher in terms of extent and durability, despite similar levels of tumor infiltration (Fig. 4C, D). We concluded that the superiority of the H1-14D is likely due to a higher affinity of the H1-14D scFv, compared to those of H1-15D or H2-14D scFvs.

**Fig. 4.**
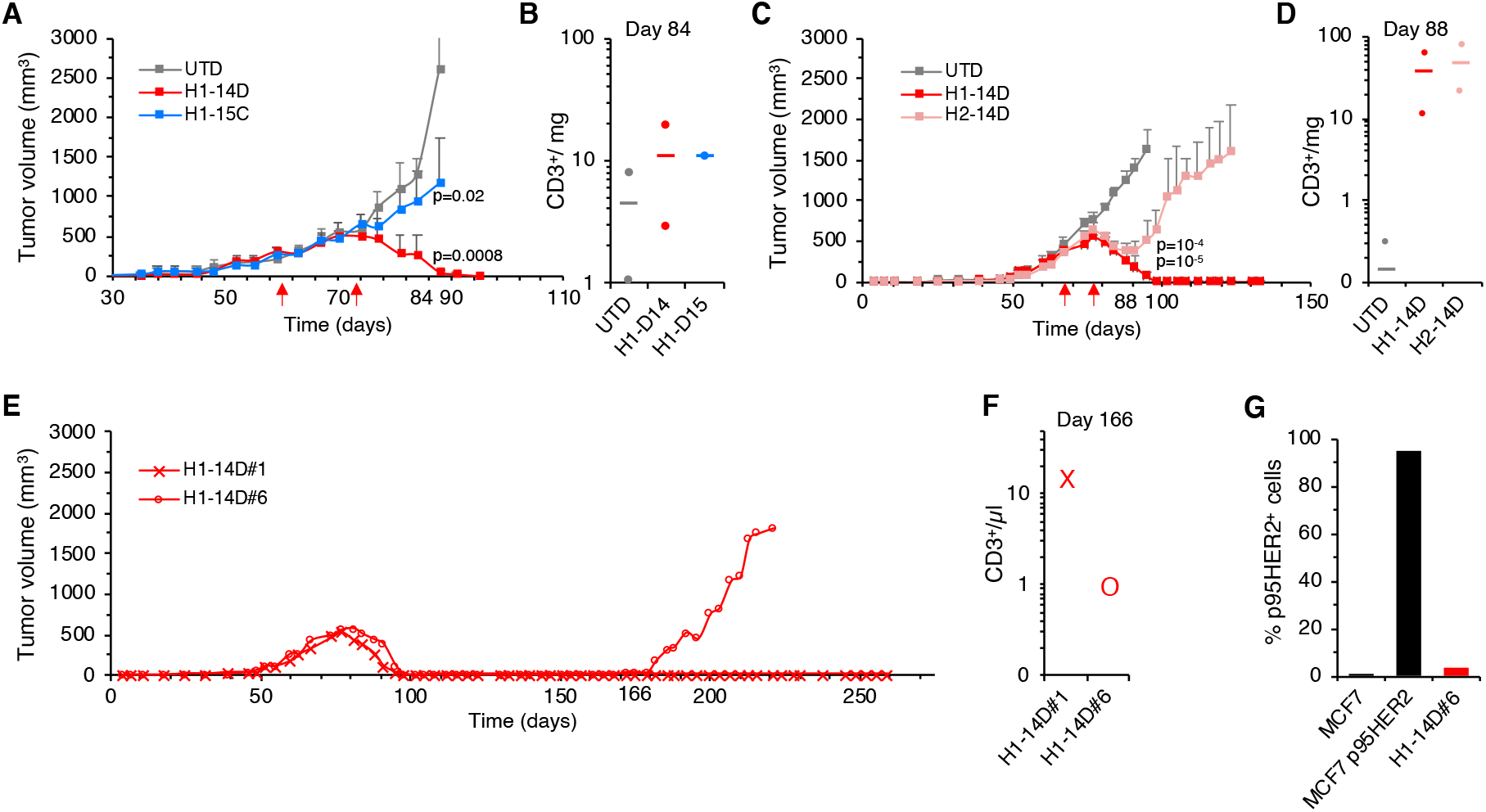
Effect of humanized p95HER2 CAR Ts *in vivo*. **A**, 3×10^6^ MCF7 cells expressing p95HER2 were orthotopically injected into NSG mice. At the time points indicated by the arrows, 3×10^6^ T cells expressing a control non-targeting CAR (UTD) or different p95HER2 CARs (H1-14D or H1-15C) were injected i.v. Tumor volumes are represented as averages ± SD (standard deviation) (n=6 per arm). **B**, Two mice from each treatment group were sacrificed at day 84. Tumors were removed, and the numbers of infiltrating lymphocytes were analyzed and expressed as number of CD3^+^ cells per milligram of tumor. Averages of the two measures are shown. **C, D**, A similar approach as in A, B was used to compare the effects of the p95HER2 CARs H1-14D and H2-14D. **E**, Two mice (numbers 1 and 6) from the group treated with the H1-14D p95HER CAR T cells were long-term followed up as in A. **F**, The numbers of circulating CD3^+^ cells were determined at the indicated time-point. **G**, At the end of the experiment, the tumor from mouse #6 was disaggregated and the levels of p95HER2 determined by flow cytometry. As controls we used parental MCF7 cells and the MCF7 cells expressing p95HER2.

Long-term monitoring of two mice treated with H1-14D, henceforth referred to as p95HER2 CAR for simplicity, showed a durable response and a recurrence (Fig. 4E). The presence of circulating human CD3^+^ cells (Fig. 4F) indicated that the clearance of p95HER2 CAR Ts was not the cause of this resistance. Confirming this possibility, analysis of tumor cells derived from the resistant tumors showed the loss of p95HER2 expression in the membrane (Fig. 4G). Thus, the p95HER2 CAR selected has a remarkably durable anti-tumor effect and the resistance observed was due to loss of p95HER2 expression.

### Effect of p95HER2 CAR Ts on metastases

Metastatic breast cancer remains an essentially incurable disease (Martínez-Sáez and Prat, 2021). Thus, the search for novel therapies against lung and brain metastases, two frequent sites to where breast cancer cells migrate, is warranted.

We injected MCF7 p95HER2 cells expressing luciferase into the tail vein of immunosuppressed mice. After one month, we detected cell growth in the lungs as shown by the increase in bioluminescence (Fig. 5A). Treatment with the p95HER2 CAR effectively reduced this metastatic growth (Fig. 5A).

**Fig. 5.**
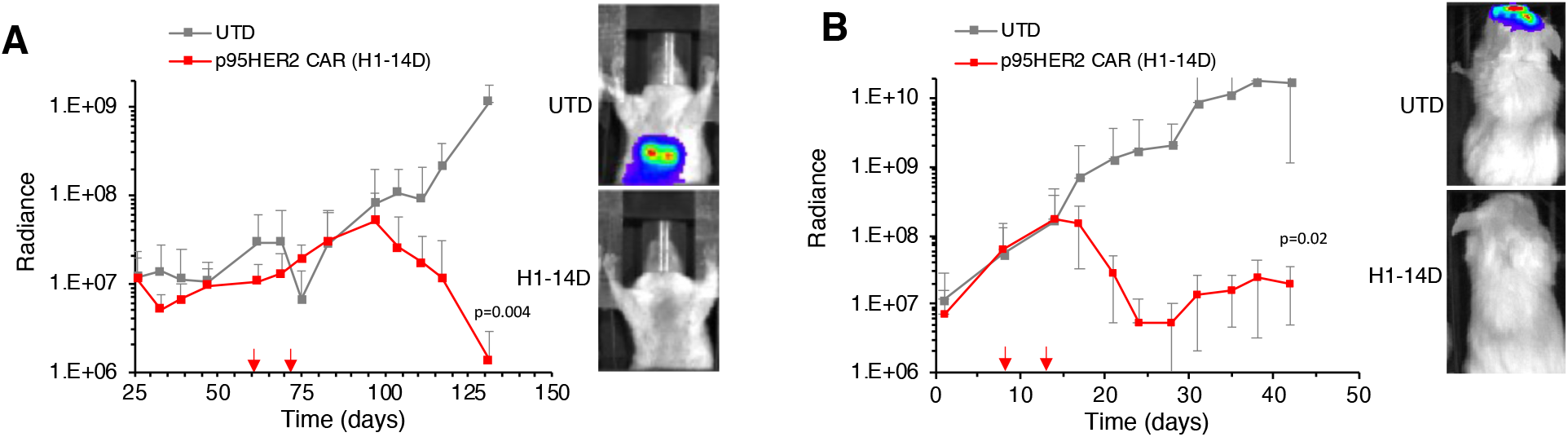
Effect of the H1-14D p95HER2 CAR T on lung and brain metastases. **A**, Left, MCF7 p95HER2 cells expressing luciferase were injected into the tail vein of NSG mice. 25 days after the injection lung metastasis were detected by *in vivo* luminescence. At the time points indicated by the arrows, 3×10^6^ CAR Ts expressing a non-targeting CAR (UTD) or the p95HER2 CAR H1-14D were injected i.v. and metastatic growth was monitored (n=3 per group). Right, representative examples of mice in each group. B, The same cells as in A were injected intracranially into NSG mice which were treated and monitored as in A (n=6 per group).

Next, we injected intracranially MCF7 p95HER2 luc cells. Soon after the injection, cells proliferated. Again, treatment with the p95HER2 CAR Ts reduced these brain metastases (Fig. 5B). We concluded that the p95HER2 CAR Ts could cross the blood brain barrier and kill p95HER2 expressing tumor cells growing in the brain.

Collectively, the results presented here show that second generation CAR Ts targeting p95HER2 are effective against cancer cells expressing p95HER2 in different tissues.

## Discussion

While most HER2-positive breast tumors diagnosed at early stages are currently being successfully treated with surgery combined with neo- and adjuvant precision therapies, advanced tumors still pose an unmet medical need. In addition, although increasingly infrequent, late recurrences of apparently eradicated tumors still occur, and they tend to not respond to currently available anti-HER2 therapies (Martínez-Sáez and Prat, 2021). Further underscoring the need for additional anti-HER2 therapies, other HER2-positive tumors, such as those affecting the gastrointestinal tract, do not respond to current therapies to the same extent as HER2-positive breast tumors do.

In this context, the redirection of T cells against HER2-positive tumor cells is an attractive therapeutic avenue. One of the difficulties to treat solid tumors with CAR T cells are the on-target off-tumor effects. This hurdle is evident in the case of HER2, which is expressed is many epithelial tumors, albeit at much lower levels than in amplified tumors. In fact, fatal side effects have been already documented in a patient treated with CAR T cells targeting HER2 (Morgan et al., 2009).

Since p95HER2 is a *bona fide* tumor specific antigen, on-target off-tumor effects are not expected, and thus, p95HER2 CAR Ts are likely a safe alternative to HER2 CAR Ts. The results shown in this report strongly support this possibility; p95HER2 CAR Ts are effective against p95HER2-expressing cells both *in vitro* and *in vivo*. The remarkable *in vivo* response observed can be explained by two hypothetical mechanisms. CAR Ts could be effective enough to wipe out all tumor cells, resulting in the durable responses documented in Fig. 4. Alternatively, CAR Ts can persist long enough to keep in check any persistent cancer cells. The detection of CAR Ts more than 150 days after administration favors the second possibility.

Loss of antigen has been described as a putative mechanism of resistance to CAR Ts (Rafiq et al., 2020). Accordingly, cell engineered to express p95HER2 can become resistant to p95HER2 CAR T by dowmodulating the former (Fig. 4). The mechanism of generation of p95HER2 in these cells - ectopic expression of a cDNA encoding p95HER2- is different from that occurring in tumors - alternative initiation of translation-. Thus, the downmodulation of p95HER2 as a mechanism of resistance should be evaluated in experimental systems closer to real tumors, such as patient-derived xenografts.

Additional mechanisms of resistance are expected to arise in tumors treated with p95HER2 CAR Ts. The main advantage of CAR Ts is that they can be used as platforms to deliver additional anti-tumor factors such as cytokines or antibodies to counteract these mechanisms. The cytokines produced by these so-called fourth generation CARs include IL-12 (Pegram et al., 2012), IL-18 (Chmielewski and Abken, 2017; Hu et al., 2017) and IL-15 (Hoyos et al., 2010). The antibodies delivered by fourth generation CARs Ts include blocking antibodies targeting immune checkpoint inhibitors (Li et al., 2017; Suarez et al., 2016) and bispecific antibodies (Choi et al., 2019; Yin et al., 2021).

In summary, p95HER2 CAR T cells represent a new avenue to treat a subset of HER2-positive tumors that, in the near future, will be complemented with additional anti-tumor therapies to achieve durable anti-tumor responses.

## Materials and Methods

### Cell lines

MCF7 (#HTB-22) and MCF10A (#CRL-10317) were purchased from American Type Culture Collection (ATCC). MCF7 & MCF10A cells expressing p95HER2 were generated as described previously (Rius Ruiz et al. 2018). Cells were maintained in Dulbecco’s Modified Eagle’s Medium:F12 (DMEM:F12, Gibco) supplemented with 10% fetal bovine serum (FBS, Gibco) 1% L-glutamine (Biowest) and the corresponding vector resistance antibiotics.

PBMCs were cultured in RPMI GlutaMAX medium (Gibco), HEPES (Sigma-Aldrich) and 1% penicillin/streptomycin (Sigma-Aldrich).

Mycoplasma contamination was tested using the kit MycoAlertTM (Lonza). Cells were not authenticated in-house.

### p95HER2 Antibodies

Murine antibodies targeting human p95HER2 were generated using hybridoma technology. Humanization of the heavy (Vh) and light (Vl) chains of the best two candidates was performed by Roche using proprietary technology. Three humanized versions of each murine antibody were produced, sequenced and tested.

Dot blot assay to determine the epitope of the antibodies was performed as follows. Overlapping 8-mer peptides from the extracellular region of human HER2 were immobilized to a nitrocellulose membrane and antibodies were incubated for 30 minutes. Membranes were developed using peroxidase-linked secondary antibody.

### Generation of CAR constructs

All CAR constructs containing the different p95HER2 scFvs were synthetized and cloned into the retroviral vector pMSGV1, which includes the murine stem cell virus (MSCV) long-terminal repeat (LTR) as promoter. The constructs contain a CD8 hinge domain with an intracellular CD28 co-stimulatory domain and a CD3z signaling domain. The complete sequences of the anti-p95HER2 scFvs are described.

### PBMCs isolation

PBMCs were isolated from fresh buffy coats obtained from healthy donors through the Blood and Tissue Bank of Catalonia (BST). Blood was diluted 1:2 with 1× phosphate-buffered saline (PBS) and transferred to a 50 ml falcon tube with Ficoll-Paque PLUS (GE Healthcare) at a 1:2 ratio, following the manufacturer’s instructions. After obtaining the buffy coat, red blood cells (RBC) were lysed with 1× RBC lysis buffer (Invitrogen) for 5 min. PBMCs were counted and frozen at −80 °C and used later for transduction of CAR constructs.

### CAR T cells production

Human T cells were activated by adding 300 IU ml^-1^ interleukin-2 and an anti-CD3 antibody (OKT3 clone) to PBMCs in their culture media and cultured for 2 days. Retroviruses corresponding to different CAR constructs were transduced into activated T cells in retronectin-coated plates (Takara). Cultures were allowed to expand for 10 days in their media supplemented with 300 IU ml^-1^ IL-2. CAR T cells were used for *in vitro* assays after day 3 or cryopreserved at liquid nitrogen at day 10 until use for *in vitro* and *in vivo* assays.

### *In vitro* assays

CAR Ts were normalized to transduction efficiency for co-culture experiments. In brief, CAR surface expression was detected with an anti-human AF647-conjugated antibody (Jackson ImmunoResearch) by flow cytometry.

For *in vitro* functional assays, 10,000 target cells were seeded in 96-well flat bottom plates and incubated at 37ºC, 5% CO_2_ overnight. CAR Ts were added the next day at different ratios of target cell_CAR+ Ts. Plates were incubated for 48 additional hours. Co-cultures were then harvested with trypsin and resuspended in 1× PBS, 2.5 mM EDTA, 1% bovine serum albumin (BSA), and 5% horse serum for flow cytometric analysis.

### Flow cytometry

For activation assays, CARs were incubated for 30 min with the following antibody mix: hCD8 and hCD25 from BioLegend. For proliferation assays, numbers of CD3-positive cells were determined as counts. For cytotoxicity assays, cell viability was determined as counts of alive EpCAM-positive cells. Samples from *in vivo* experiments were analyzed using hCD45 (Biolegend) and hCD3 (Biolegend).

The viability marker Zombie Aqua (Biolegend) was used in all assays to determine alive cells. Samples were assayed in the cytometer LSR Fortessa (BD Biosciences) and analyzed using FlowJo software.

### *In vivo* tumor growth assays

MCF7-p95HER2 cells (3×10^6^) were implanted into the fourth pad of 6-8 weeks NOD.Cg-Prkdcscid Il2rgtm1WjI/SzJ NOD scid gamma (NSG) mice (Charles River Laboratories) and beta-estradiol (Sigma-Aldrich) was added to drinking water. p95HER2 expression was induced with doxycycline (1 g/liter) in drinking water. Tumor volume was measured using calipers twice a week using the formula (length x width^2^) x (π/6) and mouse body weight was monitored once a week.

Once tumors reached 300 mm^3^, mice were treated intravenously with 3×10^6^ CAR T cells twice. Mice blood and tumors were collected and analyzed at the indicated days. Tumors were weighted, mechanically disrupted and enzymatically digested with Collagenase/Hyalurodinase in RPMI medium for 1 hour at 37ºC shaking at 80 rpm. The samples were then filtered through 100 mm strainers and red blood cells were lysed. Cells were resuspended in PBS with EDTA 2.5 mM and used for flow cytometry analyses. T cell infiltration data is represented as number of CD3-positive cells per milligram of tumor or per microliter of blood.

Animal work was performed according to protocols approved by the Ethical Committee for the Use of Experimental Animals at the Vall d’Hebron Institute of Oncology.

### Metastases assays

To mimic brain metastases, MCF7-p95HER2 cells (5×10^5^) transduced with luciferase as in (Rius Ruiz et al., 2018) were injected intracranially in NSG mice. For lung metastases, MCF7-p95HER2 luc cells were injected in the tail vein of NSG mice and after 1 month tumor cells colonized the lungs.

Mice were maintained with doxycycline treatment (1g/liter) and were treated intravenously with 3×10^6^ CAR T cells twice. Tumor growth was monitored via bioluminiscence imaging with the IVIS-200 Imaging System from Xenogen (PerkinElmer).

### Statistics

Data are represented as average ± standard deviation and were analyzed using the two-sided Student’s t-test using Excel. Statistical significance was considered with p-value less than 0.05.

In *in vivo* experiments, mice were allocated randomly in treatment groups and growth curves were compared using two-way analysis of the variance (ANOVA) with subsequent Bonferroni correction. Efforts were made to reduce the number of animals used in the experiments.

